# Evolution of satellite plasmids can stabilize the maintenance of newly acquired accessory genes in bacteria

**DOI:** 10.1101/669465

**Authors:** Xue Zhang, Daniel E. Deatherage, Hao Zheng, Stratton J. Georgoulis, Jeffrey E. Barrick

## Abstract

Plasmids play a principal role in the spread of antibiotic resistance and other traits by horizontal gene transfer in bacteria. However, newly acquired plasmids generally impose a fitness burden on a cell, and they are lost from a population rapidly if there is not selection to maintain a unique function encoded on the plasmid. Mutations that ameliorate this fitness cost can sometimes eventually stabilize a plasmid in a new host, but they typically do so by inactivating some of its novel accessory genes. In this study, we identified an additional evolutionary pathway that can prolong the maintenance of newly acquired genes encoded on a plasmid. We discovered that propagation of an RSF1010-based IncQ plasmid in *Escherichia coli* often generated ‘satellite plasmids’ with spontaneous deletions of accessory genes and genes required for plasmid replication. These smaller plasmid variants are nonautonomous genetic parasites. Their presence in a cell drives down the copy number of full-length plasmids, which reduces the burden from the accessory genes without eliminating them entirely. The evolution of satellite plasmids may be favored relative to other plasmid fates because they give a more immediate fitness advantage to a cell’s progeny and because the organization of IncQ plasmids makes them particularly prone to certain deletions during replication. Satellite plasmids also evolved in *Snodgrassella alvi* colonizing the honey bee gut, suggesting that this mechanism may broadly contribute to the importance of IncQ plasmids as agents of bacterial gene transfer in nature.

**Significance Statement:** Plasmids are multicopy DNA elements found in bacteria that replicate independently of a cell’s chromosome. The spread of plasmids carrying antibiotic-resistance genes to new bacterial pathogens is a challenge for treating life-threatening infections. Often plasmids or their accessory genes encoding unique functions are lost soon after transfer into a new cell because they impose a fitness burden. We report that a family of transmissible plasmids can rapidly evolve ‘satellite plasmids’ that replicate as genetic parasites of the original plasmid. Satellite plasmid formation reduces the burden from the newly acquired genes, which may enable them to survive intact for longer after transfer into a new cell and thereby contribute to the spread of antibiotic resistance and other traits within bacterial populations.

## Introduction

Horizontal gene transfer (HGT) is a dominant genetic mechanism for disseminating novel traits, including antibiotic resistance, among microbial species and populations (1, 2). Conjugative and broad-host-range (BHR) plasmids are especially important for the transfer of genes between phylogenetically distant bacterial species (3). Unlike the case for HGT via natural transformation or phage transduction, plasmid transfer by conjugation is not restricted by the requirement that DNA recombination and repair pathways integrate new genes into a bacterial chromosome. BHR plasmids, in particular, encode genes that carry out some of their own replication functions, which makes them less dependent on host factors than other plasmids and, therefore, more likely to successfully replicate in many types of bacterial cells. BHR plasmids can be classified into several different incompatibility groups on the basis of their replication mechanisms. Many IncP-1, IncN, IncW, and IncQ plasmids confer a wide spectrum of antibiotic resistance and are the most concerning with respect to the spread of multidrug resistance (4, 5).

Recently acquired plasmids generally impose a fitness burden on a new host cell, which may be due to the cost of expressing antibiotic resistance genes (6) or due to plasmid-encoded genes interfering with host processes (7–9). Therefore, purifying selection can rapidly lead to the loss of genes encoded on a plasmid from a new host unless there is positive selection for traits they confer (10). Continuous conjugative transfer to spread a plasmid to new cells within a population can counteract the purifying selection process to some extent, but there is also a fitness cost of increasing the conjugation rate on donor cells (11). Compensatory mutations on the chromosome and/or plasmid can sometimes ameliorate the cost of a plasmid and favor its persistence (12–17). However, it takes a long time, typically several hundred cell generations, with constant selection for novel plasmid function to evolve and fix these types of compensatory mutations in laboratory experiments (12, 18, 19). These conditions may be unrealistic in most natural settings.

Most experimental work examining BHR plasmid stability has used IncP plasmids as a model (13, 14, 17). Fewer studies have focused on IncQ plasmids, though these plasmids are found frequently in clinical and environmental settings and are known to confer a wide range of antibiotic resistance activities (20–23). IncQ plasmids contain replicon and mobilization modules in the backbone (24). The RSF1010, R1162, and R300B plasmids isolated independently from *Escherichia coli, Pseudomonas aeruginosa,* and *Salmonella enterica* serovar *Typhimurium,* respectively, are the best-characterized members of this group (4, 25, 26). In each of these plasmids, the minimal replicon consists of *repA, repB* and *repC* genes, which encode helicase, primase, and iteron-binding proteins, respectively, and an *oriV* region containing the origin of replication (27). Bidirectional replication of incQ plasmids starts from the *ssiA* and *ssiB* initiation sites in *oriV* and occurs via an unusual strand displacement mechanism. In this replication mode, only one strand of DNA is replicated at a time from a given plasmid template molecule, while the other strand is displaced and released as single-stranded DNA (24, 28).

All IncQ plasmids encode multiple mobilization genes, including *mobA, mobB, mobC, mobD, mobE,* and the transmission initiation site, *oriT* (24). During conjugative mobilization, the relaxosome nicks the DNA at the *oriT* site and drags the single-stranded DNA to the mating channel. Due to the IncQ replication and mobilization mechanisms, most of these plasmid molecules exist in cells as single-stranded DNA. For example, over 70% of plasmid DNA was found to be single-stranded by electron microscopy in R1162 plasmid preparations (29). This single-stranded nature makes the plasmid prone to recombination, and this propensity is thought to restrict the size of IncQ plasmids. Natural isolates are typically in the 5–14 kb range and not larger (30). IncQ plasmids are not self-transmissible. They rely on machinery from other conjugative plasmids (e.g., IncP or IncN plasmids) or host cells for DNA transfer (31, 32).

IncQ plasmids with nearly identical sequences have been isolated in organisms from distant geographic locations (4, 25, 26), and identical IncQ plasmids have been isolated from the same environment at times separated by more than 30 years (22). Both observations suggest that IncQ plasmid are evolutionarily stable. Bioinformatics analyses show that the accessary genes carried by conjugative plasmids are often integrated into the plasmid by processes mediated by other mobile genetic elements (MGE), such as transposons inserting into the plasmid, but that they can also be acquired directly by recombination mediated by short homologies (20, 21, 33). The fitness burden of a plasmid on a host cell typically increases rapidly with the number of antibiotic resistance genes it carries (34). The mechanisms underlying how IncQ plasmids are able to acquire and maintain these and other accessory genes are still not completely understood.

In this study, we performed experimental evolution of bacteria with a newly acquired RSF1010-derived IncQ plasmid. Satellite plasmids with deletions of accessory and replication genes rapidly arose that reduced the burden of plasmid carriage through a novel evolutionary mechanism. We characterized this process by monitoring accessory gene function with flow cytometry, by using next-generation sequencing to examine plasmid evolution, and by measuring the relative fitness of cells representing different endpoints of plasmid evolution. We also observed satellite plasmid formation in *Snodgrassella alvi* wkB2, a bacterium from the honey bee *(Apis mellifera)* gut microbiome, when it was cultured in the lab and when it colonized live bees. Finally, we examined the organization of this plasmid and performed population genetic simulations to understand how differences in mutation rates and the fitness effects of new mutant plasmids on cells and their progeny could influence the chances of satellite plasmids evolving versus alternative evolutionary pathways.

## Results

### IncQ Satellite Plasmids Rapidly Evolve in *E. coli*

We used an experimental evolution approach to directly observe bacterial adaptation following acquisition of a broad-host-range IncQ plasmid. Specifically, we added the plasmid pQGS (**Fig. 1A**), which has an RSF1010-derived backbone from pMMB67EH (35), to *E. coli* K-12 strain BW25113 (36). The constitutively expressed *gfp* gene on pQGS is an easily trackable proxy for an accessory function that is not under selection at the time that the plasmid is acquired by a new cell. Both this *gfp* gene and the adjacent *lacI* gene, which also has no function related to the survival of these cells, can be thought of as representing genes for resistance to an antibiotic or other stressor that is not yet present in the environment but could be in the future. pQGS also encodes the *aadA* spectinomycin resistance gene that we used to select for plasmid maintenance during the evolution experiment. This marker represents a gene that enables cells that acquire the plasmid to survive and flourish immediately, in their current environment. In the evolution experiment, six populations of *E. coli* BW25113 carrying pQGS (designated B1–B6) were serially transferred through 1:2000 daily dilutions into fresh medium such that they underwent ~330 total generations (cell doublings) over 28 total days of growth in liquid media.

**Fig. 1.**
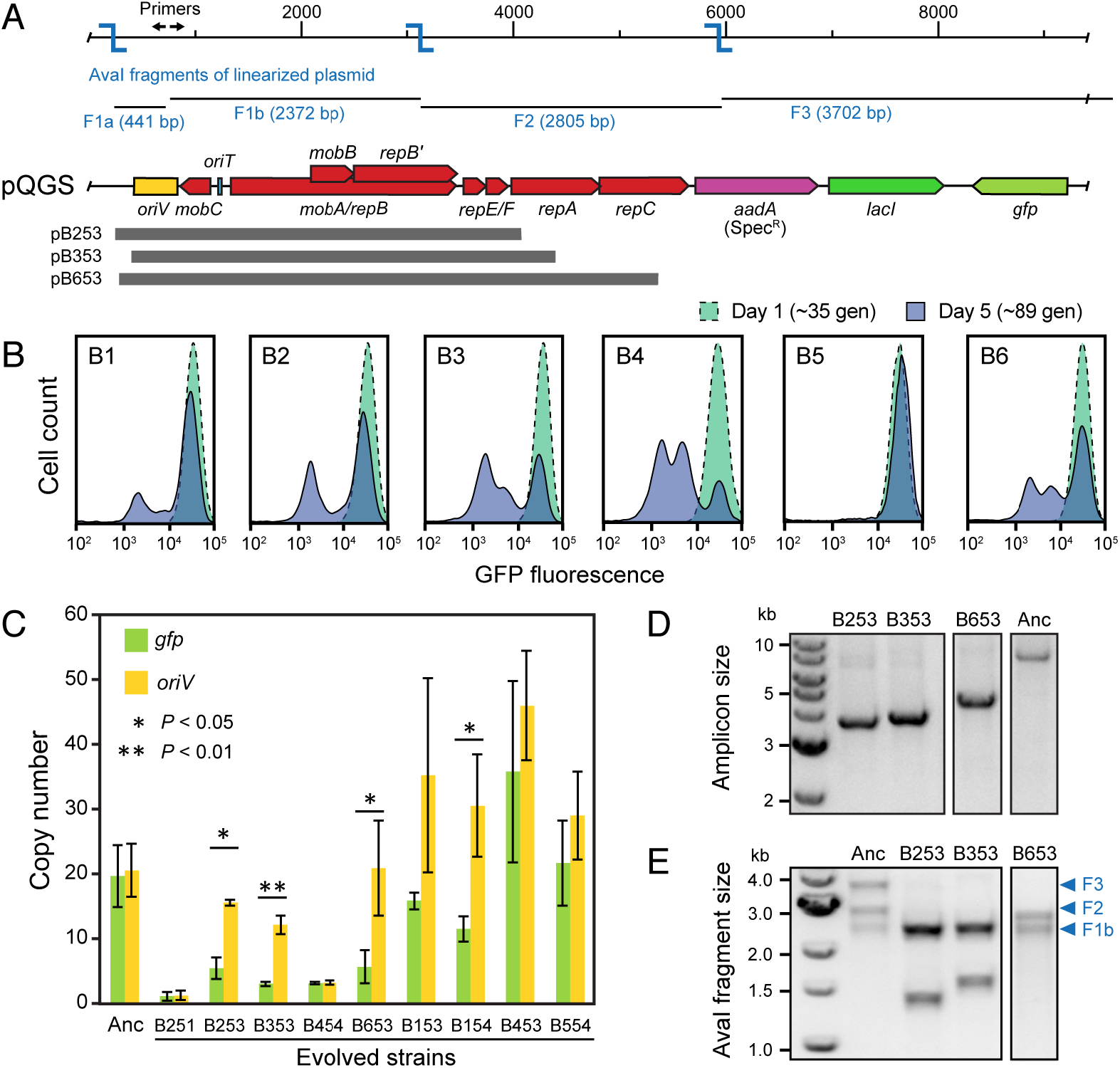
Satellite plasmids evolve from an IncQ plasmid in *E. coli.* (A) Map of the pQGS plasmid used in evolution experiments. The *aadA* gene for Spec^R^ was used to select for plasmid maintenance. Constitutively expressed *gfp* serves as a trackable proxy for a costly accessory gene. Primers used to generate PCR amplicons from pQGS for analysis (P1 and P2), AvaI restriction sites (blue), and the fragments generated by digesting the linearized plasmid with AvaI (F1a, F2a, F2, and F3) are shown above the map. Plasmid regions preserved in SPs in evolved strains B253, B353, B653 are pictured below. (B) Flow cytometry comparing the distributions of fluorescence in cell populations in all six experimental populations of *E. coli* after 1 and 5 days of evolution. (C) Copy number of the replication origin *(oriV)* and GFP gene (*gfp*) on the pQGS plasmid relative to a single-copy chromosomal reference gene (*dapA*) determined using quantitative PCR. The ancestor strain (Anc) and evolved strains from day 5 of the evolution experiment were tested. Error bars are standard deviations for three biological replicates. P values are for two-tailed t-tests comparing *oriV* and *gfp* copy number in the same strain. (D) Linearized plasmid PCR amplicons from the ancestor strain (Anc) and three evolved strains that had significantly different *oriV* and *gfp* copy numbers. (E) DNA fragments produced by an AvaI restriction digest of the samples shown in panel D. The expected fragments are labeled in panel A. The smallest fragment (F1a) does not appear on the gel. For each SP fragment, F3 is missing and F2 is truncated to an extent that is consistent with identification of the precise boundaries of the deleted regions in each SP by Sanger sequencing.

We expected mutations to arise that eliminated expression of the *gfp* accessory gene to reduce the fitness burden of the plasmid on these new host cells. Therefore, we monitored GFP levels in the populations using flow cytometry. On day 1 of the evolution experiment all six cell populations had a single uniform peak of highly fluorescent cells. By day 5, subpopulations of cells with reduced fluorescence had evolved and reached high frequencies in all but one population (**Fig. 1B**). However, we found no mutations in the *gfp* promoter or coding sequence in plasmids isolated from two random colonies with reduced fluorescence picked from each of three different populations (B2, B3, and B6). Therefore, we next tested whether these strains had evolved decreased plasmid copy number, which could also reduce *gfp* expression and plasmid burden. Using quantitative PCR, we estimated the copy numbers of the plasmid origin of replication region *(oriV)* and *gfp* gene against a single-copy chromosomal reference gene (*dapA*) (**Fig. 1C**). Some clones had equally reduced *gfp* and *oriV* copy numbers. For example, strains B251 and B454 followed this pattern (evolved clones were designated with the letter B followed by the first digit indicating the population of origin, the second digit standing for the day of isolation, and the third identifying a specific clonal isolate). This result was consistent with the evolution of a reduced plasmid sequence copy number, but it did not entirely hold for all of the evolved strains. Others we examined (B253, B353, B653, B154) had significantly different *gfp* and *oriV* copy numbers, always with more copies of *oriV* than *gfp* (**Fig. 1C**).

To figure out the reason for the discrepancy between *oriV* and *gfp* copy numbers in these strains, we further characterized plasmids isolated from three representative clones (B253, B353, B653). First, we checked plasmid size by electrophoresis. The IncQ plasmid is difficult to molecularly characterize due to its low copy number and the high prevalence of single-stranded DNA resulting from its strand displacement replication mechanism (37). Therefore, we used PCR with outward facing primers located inside *oriV* to create linearized amplicons for visualization (**Fig. 1A**). Compared to the ancestor, strains with different *oriV* and *gfp* copy numbers contained smaller plasmids (from 3 kb to 5 kb) (**Fig. 1D**). The PCR amplicons were purified and digested with the AvaI restriction enzyme to determine roughly which regions of the plasmid were missing in these strains. For the ancestral plasmid, AvaI digestion of the linearized plasmid yields four fragments, one of which (F1a) is too small to visualize (**Fig. 1A**). Compared to the ancestral plasmid, all three of these evolved plasmids kept the *oriV* bearing fragment (F1b), but the other two visible fragments (F2, F3) were either truncated or lost (**Fig. 1E**).

Sequencing these PCR products allowed us to precisely localize the missing plasmid pieces. A region beginning within *repA* or *repC*, including the *gfp* and *aadA* genes, and ending before *oriV* was deleted from each plasmid (**Fig. 1A**). The *repA* and *repC* genes encode the DNA helicase and iteron-binding protein, respectively, which are required for initiation of plasmid replication (27). Loss of either function should lead to a plasmid that is unable to replicate on its own. As expected, attempts to transform the shorter plasmid variants into new *E. coli* cells were unsuccessful. The PCR assay will preferentially amplify these smaller plasmids when they are present in a mixture, but weak bands corresponding to the linearized full-length pQGS plasmid are visible in the same sample in some cases (**Fig. 1D**). Since the qPCR results also show that copies of the *gfp* gene are still present in all three of these strains (**Fig. 1C**), we conclude that the smaller plasmids are satellite plasmids that co-exist with and require copies of the full-length plasmid in the same cell. From the perspective of the original plasmid, these satellite plasmids are parasites. They take advantage of its replication machinery without encoding all of the components needed for autonomous replication or any of the accessory genes.

We next examined how prevalent satellite plasmid (SP) evolution was in our experiment. Using the PCR assay, we detected SPs in all 6 populations on day 5 (**Fig. S1**). SP formation was so widespread that multiple different SPs were even observed in the same population in two cases (B1 and B4). Sequencing the linearized PCR products from clonal isolates carrying each of these new SPs revealed that they were similar to the initially characterized plasmids. All had deletions that completely eliminated the *gfp* and *aadA* genes, overlapped one or more *rep* genes, and left the *oriV* sequence intact (**Fig. S2**). The deletions leading to SP formation were nearly always flanked by short, near-perfect sequence homologies with lengths of 7 to 15 base pairs (**Table S1**), suggesting that SPs may form through RecA-independent processes (38).

### Accessory Genes are Eventually Lost or Integrated into the Chromosome

To understand the role of satellite plasmids in the maintenance of accessory genes, we further examined the time courses of plasmid evolution in the experimental populations. We found plasmids with deletions leading to 3.5–6.0 kb PCR amplicons had evolved in all six populations by day 5, persisted through day 9 in all but one population, and were still present on day 15 in three of the six populations (**Fig. 2A, B** and **Fig. S3A**). Sequencing these PCR products showed that they were all nonautonomous SPs with deletions overlapping replication and accessory genes (**Fig. S1**). In four populations (B3, B4, B5, B6), larger PCR amplicons with sizes of 7.0–8.0 kb emerged at later time points as satellite plasmids disappeared. These amplicons corresponded to various plasmids with accessory gene deletions that included loss of the entirety of *lacI* gene and a portion of *gfp* in every case (**Fig. S2**). Complete sets of replication proteins and the *aadA* gene were retained in these plasmids, meaning that they remained fully autonomous. In the other two populations (B1 and B2), SPs disappeared, and no plasmid amplicon of any type was detectable on day 21 and day 25 (**Fig. 2A** and **Fig. S3A**).

**Fig. 2.**
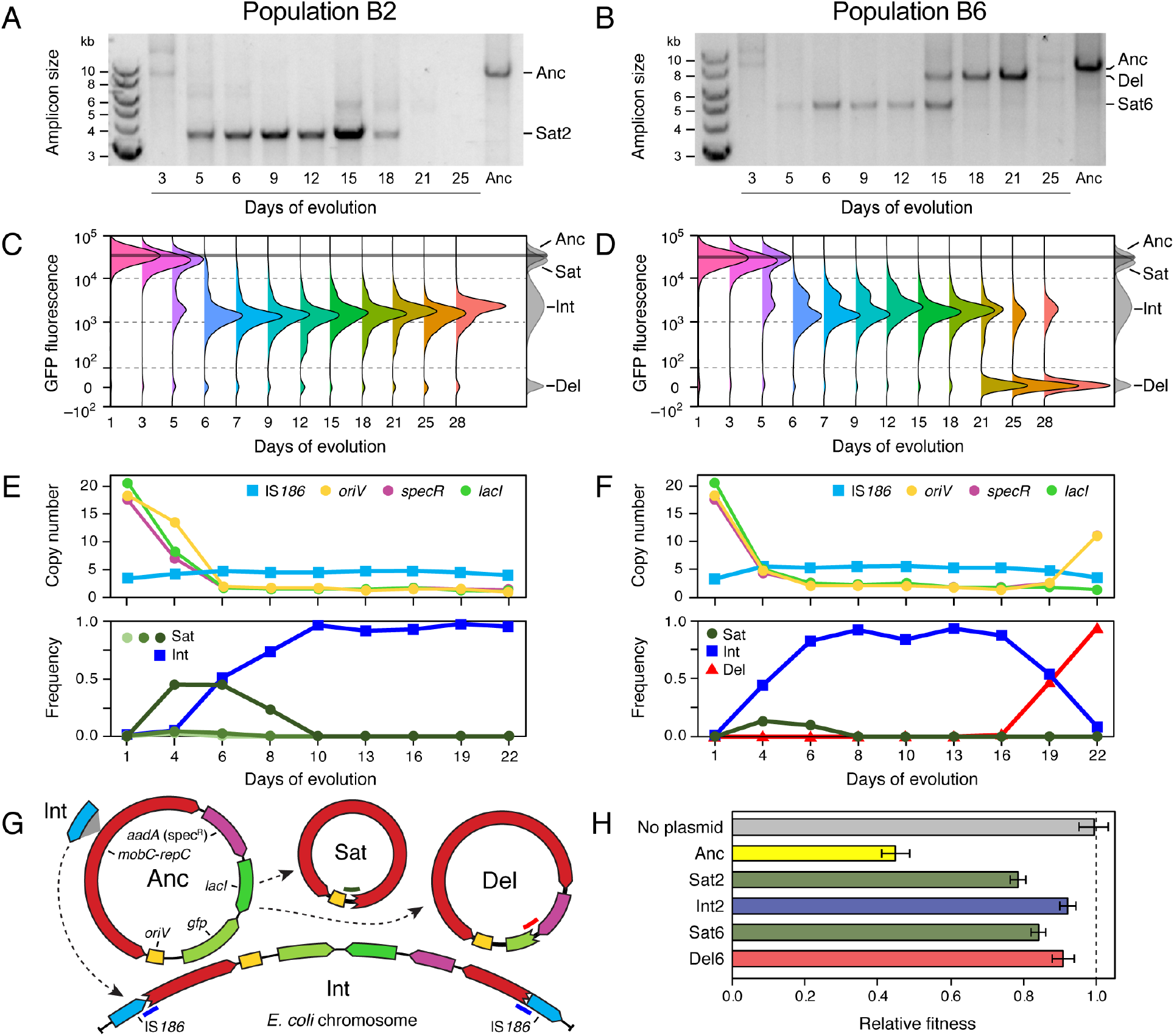
Time courses of plasmid evolution in *E. coli.* (A, B) Appearance and persistence in populations B2 and B6 of satellite plasmids and other evolved plasmids with deletions that also change the size of the linearized plasmid PCR amplicon. (C, D) Distributions of GFP expression in cells in each population determined using flow cytometry. (E, F) Estimates of the copy numbers of different plasmid regions and the relative frequencies of different evolved plasmids fates in each population from Illumina sequencing data. (G) Schematic showing different evolved fates of the pQGS plasmid. Bold lines in blue, red, and dark green show the locations of sequencing reads unique to integrated, deletion and satellite, plasmids, respectively, which were used to estimate their relative frequencies. (H) Relative fitness of strains reconstructed with different evolved plasmid variants determined using co-culture competition assays versus a control strain with no plasmid.

For two representative populations of each type (B2 and B6), we correlated flow cytometry data (**Fig. 2C, D**) with whole-genome sequencing (WGS) data (**Fig. 2E, F**) to characterize plasmid evolution in a way that was not subject to PCR amplification biases. In addition to better defining evolutionary dynamics involving satellite and deletion plasmids, these data revealed an additional plasmid fate (**Fig. 2G**). In both populations, WGS reads consistent with an insertion of an IS*186* element into the pQGS plasmid *repF* gene were detected by day 6 (**Fig. 2E, F**). IS*186* is one of the most active insertion sequences in *E. coli.* It inserted into pQGS at a site matching its preferred target site of 5’-GGGG(N6/N7)CCCC (39–41). This IS insertion likely creates a second class of satellite plasmid that is unable to replicate autonomously because it separates the *repA* and *repC* genes from their promoter which is located upstream of *repE* gene (27).

At the same time we detected the IS*186* insertions, there was a precipitous drop in the copy number of the plasmid origin of replication *(oriV)* in both populations (**Fig. 2E, F**), and subpopulations of cells with greatly reduced fluorescence began to dominate (**Fig. 2C, D**). These results are consistent with integration of the IS*186*-disrupted plasmid sequences into the *E. coli* chromosome, either concomitant with IS*186* insertion or shortly thereafter, through single-crossover recombination with one of the three original copies of IS*186* (**Fig. 2G**). We were able to detect hybrid PCR products when using one chromosomal and one plasmid primer, indicating that this was the case. The presence of multiple peaks with intermediate fluorescence intensities—at the same time in population B6 (**Fig. 2D**) and later and to a lesser extent in population B2 (**Fig. 2C**)—might indicate that the integrated plasmid sequence is sometimes duplicated within the chromosome. This could occur through homologous recombination during chromosomal replication, facilitated by the flanking IS*186* copies (**Fig. 2G**).

We also tracked the prevalence of satellite plasmids from the numbers of WGS reads that spanned deleted regions. SPs first appeared by day 4 in both populations (**Fig. 2E, F**). In population B2, we could detect three different sizes of SPs. SP incidence as a percentage of all plasmids in the population peaked at ~50% in B2 and ~20% in B6. In population B2, the copy number estimated from the read-depth coverage of the accessory gene completely deleted in all

SPs *(lacI)* dropped noticeably more quickly than that of the origin *(oriV),* as expected if satellite plasmids and ancestral plasmids coexisted in a sizable number of cells in the population. SPs dropped below the detection limit by WGS at day 9 and 7 in populations B2 and B6, respectively. At this point the IS-mediated integrants dominated. Completely nonfluorescent cells became dominant in population B6 near the end of the experiment (**Fig. 2D**). A new plasmid variant with a deletion of *lacI* and a portion of *gfp* was detected at day 19 by WGS in this population (**Fig. 2F**). According to the flow cytometry data, the prevalence of cells containing only the deletion plasmid increased to 47% at day 21 and 66% at day 25.

Overall, these evolutionary trajectories suggest that satellite plasmid evolution was beneficial to the fitness of a cell because it curtailed the expression of *gfp* and/or *lacI.* These results further suggest that IS*186*-mediated integration of the plasmid into the chromosome or deletion of these accessory genes from an evolved plasmid that remained capable of self-replication conferred greater fitness benefits than SP formation. Flow cytometry time courses show that these two fates eventually dominated in the other four populations of the evolution experiment (**Fig. S3B**).

To directly test the fitness consequences of evolving the different plasmid fates, we reconstructed cells of each type and competed them against a reference strain of *E. coli* BW25113 with the spectinomycin resistance gene *(aadA)* integrated into its chromosome and a restored ability to utilize arabinose as a neutral genetic marker. First, we isolated satellite plasmids from each population (Sat2 and Sat6) and retransformed them into cells harboring the ancestral plasmid. Next, we isolated the deletion plasmid from population B6 (Del6) and transformed it into the ancestral strain with no plasmid. Finally, we used transduction to move an integrated copy of the plasmid from an evolved cell in population B2 to the ancestral strain (Int2) so that its fitness could be measured without interference from any other evolved alleles.

As a control, we competed the plasmid-free ancestor strain BW25113 against the reference strain in media that did not contain any antibiotic. The fitness of the ancestral strain (No plasmid) was not significantly different from that of the reference strain (*p* = 0.768, two-tailed t-test). The ancestral strain/plasmid combination of BW25113/pQGS (Anc) had a greatly reduced fitness of 0.431 relative to the reference strain with no plasmid. The addition of a satellite plasmid to a cell significantly alleviated this cost, with a relative fitness of 0.781 and 0.840 for Sat2 and Sat6, respectively (**Fig. 2H**). The cases of integration into the chromosome (Int2) or deletion of the accessory genes from the plasmid that we tested (Del6) restored fitness even more toward that of cells with no plasmid, to relative fitnesses of 0.923 and 0.910, respectively. As predicted by the coexistence of cells with deletion plasmids and cells with chromosomally integrated plasmids in population B6, the fitness values of these evolved plasmid fates were not significantly different (*p* = 0.474, two tailed t-test). All strains with evolved plasmid variants were significantly more fit than the ancestor with pQGS (*p* ≤ 3.6’ 10^-14^, Bonferroni-corrected two-tailed t-tests) and less fit than the reference strain with no plasmid (*p* ≤ 0.011, Bonferroni corrected two-tailed *t*-tests).

### Satellite Plasmids Evolve in the Honey Bee Gut

We next used *Snodgrassella alvi* wkB2, a core gut symbiont of honeybees *(Apis mellifera)* to test whether SP evolution from plasmid pQGS would occur in a different bacterial species and in a complex environment associated with an animal host. *S. alvi* is a β-proteobacterium that has only recently been cultured (42). We first used the PCR assay to check for the evolution of SPs from pQGS within five populations of *S. alvi* wkB2 (designated S1–S5), during serial passage *in vitro* (**Fig. 3A**). By sequencing the PCR products, we determined that two identical bands detected at day 8 in all populations (pS621 and pS521) corresponded to SPs (**Fig. S2**). Bands that likely correspond to smaller SPs (e.g., pS631) became dominant by day 12 in all populations. Larger amplicons that may be accessory-gene deletion plasmids were also observed in two populations at this point.

**Fig. 3.**
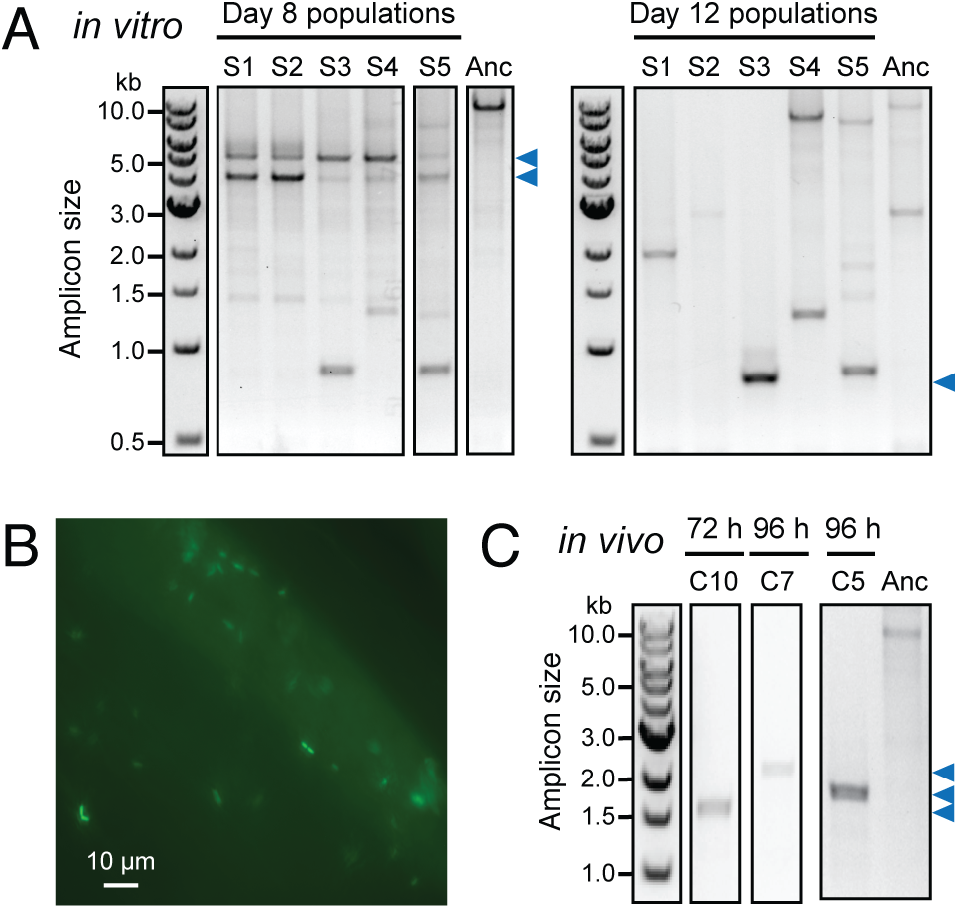
Satellite plasmids evolve in *S. alvi* in the honey bee gut. (A) SPs detected as linearized PCR amplicons during the *in vitro* propagation of *S. alvi* at days 8 and 12 in five experimental populations S1–S5. (B) Fluorescence micrograph showing colonization of the bee gut ileum 24 h after inoculation with *S. alvi* carrying plasmid pQGS. (C) Satellite plasmids detected as linearized PCR amplicons from DNA extracts of the guts of individual bees reared in different enclosures (C1–C10) at 72 or 96 h after inoculation. Full results for all populations evolved *in vivo* are shown in **Figure S4**. Blue arrows in all panels are bands that were Sanger sequenced to validate that they represent SPs (**Fig. S2** and **Table S1**).

Next, we examined whether SPs would evolve and persist in *S. alvi* wkB2 carrying pQGS when they colonized microbiota-free bees (43, 44). As expected from previous studies (45), *S. alvi* with the pQGS plasmid began to colonize the inner wall of the ileum by 24 h postinoculation as visualized by GFP fluorescence in the bee gut (**Fig. 3B**). We further confirmed colonization by measuring colony-forming units in guts isolated from sacrificed bees. On day 3 and day 4, there were approximately 10^7^ and 10^8^ CFUs per bee gut, respectively. Previous studies have showed that *S. alvi* fully colonizes microbiota-free bees by 4 to 6 days post-inoculation (46). Therefore, we used PCR to check for SP formation in bees sacrificed every 24 h during the first 4 days of colonization from ten different enclosures (**Fig. 3C, Fig. S4**). Three SPs (pC562, pC765, pC1025) were identified from bees reared in different enclosures 3 to 4 days after they were inoculated. No SP was detected in bees sacrificed 8 and 12 days after inoculation, suggesting that SPs may be a transient evolutionary intermediate in *S. alvi* colonizing the honey bee gut as they were in *E. coli* and *S. alvi* passaged *in vitro.*

### Multiple Factors Favor Satellite Plasmid Evolution Over Accessory Gene Deletion

Mutations that eliminate different regions of the pQGS plasmid can result in nonautonomous satellite plasmids or in autonomous plasmids that have deleted just the accessory genes. We found that *E. coli* cells with accessory-gene deletion plasmids (DPs) have a much higher fitness than cells containing a mixture of satellite and ancestor plasmids. If both types of cells arise at similar rates in a bacterial population, then cells with DPs are expected to dominate to such an extent that SPs should never reach an observable frequency, but this was not the case in our experiments. To investigate this discrepancy, we examined ways in which the genetic organization of the pQGS plasmid and its multicopy nature might favor satellite plasmid evolution and thereby maintenance of the newly acquired accessory genes in a population.

First, there might be a mutational bias favoring the evolution of satellite plasmids. DPs must maintain the entire plasmid from *oriV* to *aadA* intact so that they remain autonomous and still encode antibiotic resistance. If we also assume that they must delete at least a portion of the both the *lacI* and *gfp* coding sequences to realize their full fitness advantage, then this constrains these deletions to relatively few combinations of starting and ending base positions. On the other hand, SPs must only maintain *oriV* while deleting the accessory genes plus *aadA* and least one of the replication genes. Given just the raw numbers of start-end coordinate combinations fitting each of these scenarios (**Fig. 4A**), SPs would be expected to arise at 4.0 times the rate of DPs.

**Fig. 4.**
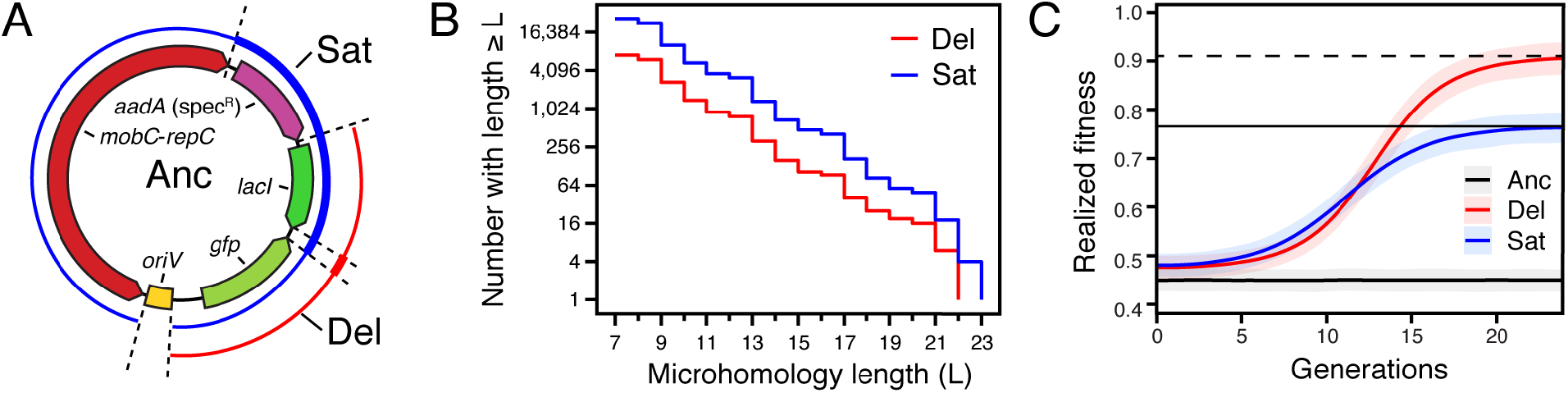
Satellite plasmid evolution is favored over deletion plasmid evolution. (A) pQGS plasmid map showing regions containing the endpoints of deletions leading to satellite plasmids (blue thin line) and accessory-gene deletion plasmids (red thin line). The regions required to be contained in each deletion are shown with bold lines in the respective colors. (B) Microhomology regions like those observed flanking satellite plasmid deletions are more common in the satellite plasmid deletion endpoint regions than in the accessory-gene deletion plasmid endpoint regions, but the difference is no greater than the 4-fold difference expected if these sites are randomly distributed. (C) Multilevel stochastic simulations predict an early advantage and due to a reduced phenotypic lag for cells with newly evolved satellite plasmids and their progeny realizing their full fitness benefit compared to deletion plasmids. Shaded areas are 95% confidence intervals from 200 simulation trajectories.

All ten unique SPs we characterized resulted from deletions that had DNA sequence microhomologies consisting of near-exact 7 to 15 base-pair repeats overlapping their endpoints in the ancestral plasmid (**Table S1**). We investigated whether there was any bias in the abundance of similar microhomologies in different regions of the plasmid that might additionally favor deletions leading to SPs versus DPs (see **Materials and Methods**). Over a range of different microhomology lengths, the number of near-repeats that could mediate the formation of each type of evolved plasmid did not noticeably deviate from the ratio of 4:1 expected if they were randomly distributed in the plasmid (**Fig. 4B**). Surprisingly, all four DPs did not have any microhomology near their endpoints (**Table S1**), despite the fact that near-exact repeats exist in these regions. This striking contrast may indicate that the mechanisms of deletions in different regions of the pQGS plasmid may vary due to how IncQ plasmids replicate (see **Discussion**).

Mutant plasmids start as a single copy within one cell, so the dynamics of plasmid replication and segregation affect how much that cell and its progeny benefit from the mutant plasmid and the chances that copies of the mutant plasmid will survive in the cell population (47). The lag between when a mutant plasmid first appears in a cell and when that cell’s descendants realize the full fitness benefit possible from harboring multiple copies of that plasmid is known as its phenotypic delay (48). We examined phenotypic delay using multilevel stochastic simulations with parameters fit to be consistent with our *E. coli* evolution experiment (see **Materials and Methods**). If SPs replicated more quickly than other plasmid types due to their smaller size, then they would be expected to have a much shorter phenotypic delay. However, models in which we gave SPs an advantage for within-cell replication were not consistent with the experimentally observed numbers of ancestral and satellite plasmids co-existing within cells in our experiment and/or the relative fitness of different cell types. Therefore, we gave all plasmids equal rates (chances) of replication in our simulations, which is consistent with a model of plasmid replication in which initiation, rather than elongation or other steps, is rate-limiting.

These simulations predict that SP cells will initially fare better in terms of their fitness before DP cells surpass them (**Fig. 4C**). Because a one-to-one trade of a copy of the ancestral plasmid for a SP is more beneficial than replacing it with a DP, the model predicts that if a cell with a SP and a cell with a DP evolved at the same time, the descendants of the SP cell would be more fit, on average, than those of the DP cell for the first 12 generations. This reduced phenotypic delay is expected to give a newly evolved SP a better chance of avoiding stochastic loss. For example, with the daily 1:2000 serial transfer regime of our *E. coli* experiment, the simulations predict that a new SP has a 17% greater chance than a new DP of establishing a population of descendant cells which is large enough that it no longer risks being randomly lost to dilution during transfer.

## Discussion

Acquisition of new antibiotic resistance genes often imposes a fitness burden on a bacterial cell. Thus, purifying selection within a bacterial population favors eliminating these and other types of newly acquired accessory genes during times when they are not important for survival or fitness (e.g., when no antibiotic is present). IncQ plasmids have frequently been identified as mobile carriers of antibiotic resistance genes in clinical environments (20–23). We found that nonautonomous satellite plasmids (SPs) that delete both accessory genes and genes necessary for plasmid replication frequently evolve within bacterial populations harboring an IncQ plasmid, both in culture and in an animal host. Because SPs are molecular parasites that reduce the copy number of full-length plasmids in a cell, SP evolution can rapidly alleviate much of the burden of newly acquired accessory genes without leading to their complete loss from a cell. Therefore, the propensity of IncQ plasmids for generating SPs can prolong the maintenance of antibiotic resistance genes in a population after it colonizes a new bacterial host.

Why are IncQ plasmids so prone to satellite plasmid formation? Nonautonomous ‘cheater’ plasmids like these can only evolve readily if a plasmid encodes factors needed for its replication separately from origin sequences that are sufficient to direct replication. In the pQGS plasmid used in this study, the origin of replication *(oriV)* is located at one end of the plasmid backbone, which makes it possible for a single deletion to remove the costly accessory genes and one or more genes encoding replication factors while leaving the origin intact. Although there are some plasmids with replication mechanisms that are independent of any plasmid-encoded proteins (e.g., ColE1 plasmids), most bacterial plasmids do encode at least one protein (usually named Rep) that is needed to initiate replication outside of the origin sequences to which it binds. Thus, these plasmids could theoretically evolve or be engineered into satellite plasmids, as long as SPs and intact full-length plasmids are able to stably co-exist within cells and their progeny.

IncQ plasmids replicate by a strand displacement mechanism and do not encode a partitioning system, so they randomly segregate into daughter cells (24). RepC has been reported to positively regulate copy number (49), while MobC and MobA suppress it (50). Despite these regulatory feedbacks, we found that total plasmid copy number was largely unchanged in cells that evolved SPs that inactivated *repC* in the *E. coli* experiments. We also observed SPs with a large range of sizes that deleted a variety of combinations of replication and mobilization genes, indicating that many different deletions are compatible with stable maintenance of these satellite plasmids. Our modeling indicates that smaller pQGS-derived SPs do not have a significant advantage over full-length plasmids in terms of their replication rate within cells. This could be another reason that it is possible for a cell to maintain enough full-length plasmids such that there is not too great a fitness cost for SP evolution due to some of a cell’s progeny inheriting only nonautonomous plasmids and thereby losing plasmid function entirely. It is possible that plasmids from other families, which commonly utilize theta or rolling-circle replication mechanisms, are subject to regulatory controls that make the presence of derived satellite plasmids more disruptive to their continued replication and segregation.

The rate at which new satellite plasmids are generated during plasmid replication is another important factor in whether they will be observed relative to other plasmid fates. We investigated whether there were aspects of SP organization and population dynamics that could favor their evolution in our experiments, especially relative to plasmids that delete burdensome accessory genes but maintain the ability to self-replicate. We found that there are more opportunities for deletions that generate SPs (i.e., mutations that generate them are expected to be more common) and that they are expected to initially give greater fitness benefits to the host cell and its progeny (i.e., they have a reduced phenotypic delay). However, their rather slight advantages in these areas are not enough on their own to explain the prevalence of SPs in our experiments.

Our observation that the deletions which create satellite plasmids are flanked by sequence microhomologies but those that generate accessory-gene deletion plasmids are not, suggests that the unusual strand displacement mechanism of IncQ plasmid replication may be a key reason that SPs evolve in our experiments. This mechanism begins with RepC binding to the iteron in *oriV* and disrupting the DNA double helix. Then, RepA unwinds the DNA and exposes the *ssiA* and *ssiB* single-strand initiation sites to the primase RepB. RepB primes DNA synthesis in a 5’ to 3’ direction separately on both strands producing large quantities of single-stranded plasmid DNA (ssDNA) intermediates (24, 51). The mobilization mechanism of IncQ and other plasmids, in which double-stranded plasmid molecules are nicked by the plasmid-encoded relaxase at *oriT* and unwound for conjugative transmission, further contributes to the generation of large amounts of single-stranded plasmid DNA inside of a host cell (27, 52, 53). ssDNA is key player for RecA-independent recombination mediated by annealing of short sequence homologies (54–56). We hypothesize that the presence of large amounts of ssDNA in cells with the pQGS plasmid may explain why SPs evolve so quickly. Deletions that remove only the accessory genes may occur at a much lower rate and through alternative mechanisms because both of their endpoints are closer together and located on the same arm of the plasmid relative to *oriV.* Similarly, plasmids with theta or rolling-circle replication mechanisms may only rarely evolve SPs.

Formation of SPs is one mechanism that can lead to persistence of accessory genes after they are transferred to a new bacterial host. Another is that the plasmid can become temporarily or permanently single-copy and nonautonomous by integrating into the bacterial chromosome. Integration was a common endpoint in our *E. coli* evolution experiment. In the case that we fully characterized, this process occurred via insertion of an IS*186* copy into the plasmid backbone and recombination with a copy of this transposable element in the genome. Similar integration events have also been observed in evolution experiments with a different family of conjugative plasmids in *Pseudomonas putida* (10). In another instance, reversible chromosomal integration mediated by IS elements was reported for the virulence plasmid pINV in *Shigella* (57).

We have shown that a transmissible plasmid possesses an organization and replication mechanism that can favor the rapid evolution of satellite plasmids that reduce the copy number of any costly cargo genes that it has spread to a new cell. Awareness of the propensity of IncQ plasmids to evolve SPs is an important consideration when studying the ecological range of this broad-host-range plasmid and when using it to engineer cells. In this latter context, SP evolution could be interpreted as a fortuitous “safety valve” for titrating down toxic or burdensome gene expression or as an unwanted complication that leads to a different effective copy number for engineered DNA sequences than was intended (58). In summary, our results suggest that satellite plasmids may be a common evolutionary intermediate that can stabilize the spread of plasmid-borne accessory genes, including those responsible for antibiotic resistance.

## Materials and Methods

### *E. coli* evolution experiment

Plasmid pQGS was constructed from a previously described kit of genetic parts for Golden Gate assembly (58). Briefly, the RSF1010-derived pMMB67EH plasmid (ATCC 37622) backbone was assembled with the *gfp* optim-1 variant driven by the constitutive PA3 promoter. A spectinomycin resistance (*<u>aadA</u>*) marker was included to enable selection for plasmid maintenance. The full sequence of pQGS is available in Genbank (MH423581). Six individual colonies of *E. coli* BW25113 carrying pQGS were isolated to initiate the evolution experiment. Populations were grown in 10 ml of lysogeny broth (LB) (10 g NaCl, 10 g tryptone, 5 g yeast extract per liter) with 60 μg/ml spectinomycin at 37°C in 50 ml Erlenmeyer flasks with 200 r.p.m. orbital shaking over a 1-inch diameter unless otherwise noted. Cultures were transferred via a 1:2000 dilution into 10 ml of fresh LB-Spec every day. Samples of each population were periodically archived by adding glycerol as a cryoprotectant and storing at −80°C.

### Flow cytometry

Fluorescence was monitored by flow cytometry using a BD LSRFortessa instrument. Cells were counter stained by adding the red membrane dye FM 4-64 (ThermoFisher) to a final concentration of 10 ng/μl to 100 μl of cell culture. The culture was incubated for 10 min at 37°C in 1.5 ml microcentrifuge tubes with 200 r.p.m. shaking. Then, cells were spun down and stored on ice until sample loading. Right before loading, each cell sample was resuspended by adding 1 ml saline and pipetting up and down thoroughly. An aliquot of resuspended cells was diluted 1:20 with saline in 5 ml Falcon round-bottom culture tubes for loading on the flow cytometer.

### Quantification of *gfp* and *oriV* copy number by qPCR

Copy numbers of the *oriV* region and *gfp* gene on the plasmid relative to a chromosomal gene were determined by qPCR. The chromosomal gene *dapA* was set up as the reference gene (*dapA*-F: 5’-AGCGTCATCATCACCACATC, dapA-R: 5’-GTACTTCGGCGATCGTTTCT) to compare to the *gfp* gene *(gfp-F:* GAGGATGGAAGCGTTCAACTA, *gfp-R:* 5’-GCAGATTGAGTGGACAGGTAA) and *oriV* region (oriV-F: 5’-TCCAGCGTATTTCTGCGG; oriV-R: 5’-GGATAGCTGGTCTATTCGCTG) sequences on the plasmid. The *dapA* standard curve was constructed using genomic DNA isolated from BW25113 using the PureLink Genomic DNA Mini Kit (Invitrogen). Standard curves for *gfp* and *oriV* were constructed using plasmid isolated from the ancestral strain using the PureLink Plasmid DNA Miniprep Kit (Invitrogen). Total DNA amounts were quantified using a Qubit Fluorometer (ThermoFisher).

For each strain of interest, three individual colonies were isolated from revived freezer stocks and grown in 3 ml of LB-Spec in test tubes for 24 h. The cultures were then diluted 1:100 into 10 ml of fresh medium in 50 ml flasks. The cells were harvested at an OD600 of ~0.3 and genomic DNA was isolated using the PureLink Genomic DNA Mini Kit (Invitrogen). qPCR was performed using SYBR Green Real Time PCR Master Mix (ThermoFisher) at a final DNA concentration of 0.4 ng/μl using the ViiA 7 Real Time PCR System (ThermoFisher). Amplification conditions were as follows: initial denaturation for 10 min at 95°C, followed by 40 cycles of denaturation for 15 s at 95°C then annealing and extension for 1 min at 60°C.

### Detection of satellite plasmids and GFP mutations

Satellite plasmids arising in evolved populations were detected by PCR using primers P1 (5’-CCGCAGAAATACGCTGGA) and P2 (5’-CAGCGAATAGACCAGCTATCC). Purified amplicons were Sanger sequenced using the primer facing the *gfp* direction. The *gfp* gene was amplified using primers PA3-F (5’-CGGATGACACGAACTCACGA) and GFP-R (5’-GGTTCGTAACATCTCTGTAACTGCT). PCR products were purified and Sanger sequenced using both GFP primers to check mutations in the *gfp* gene and promoter region.

### Whole genome sequencing

Frozen cultures sampled during the evolution experiment were inoculated via a 1:2000 dilution into 10 ml fresh LB medium and cultured for 24 h. DNA was extracted from 1 ml of each resulting culture using the Invitrogen PureLink Genomic DNA Mini Kit (Invitrogen). This DNA was fragmented using Fragmentase (NEB). Then, Illumina sequencing libraries were prepared using the KAPA Low Throughput Library Preparation kit (Roche) according to manufacturer’s instructions, except that all reactions were carried out in one half recommended reaction volumes and both sequencing adapters were modified to include six base barcodes at the beginning of each read. These libraries were sequenced on a MiSeq instrument to generate 150-base paired-end reads at the University of Texas at Austin Genome Sequencing and Analysis Facility. Mutations were predicted by using the *breseq* pipeline (version 0.33.2) in polymorphism mode (59, 60) to compare sequencing reads to the *E. coli* BW25113 genome (GenBank: CP009273.1) and pQGS plasmid (Genbank:MH423581). FASTQ read files are available from the NCBI Sequence Read Archive (SRP149032).

### Competitive fitness assays

BW25113 is unable to utilize arabinose (Ara^−^) due to the *Δ(araD-araB)567* mutation. We created an Ara+ variant of BW25113 as a reference strain to compete against Ara^−^ strains from the evolution experiment. To do so, we first inserted a Spec^R^ marker upstream of the intact *ara* operon in *E. coli* strain MG1655 using Lambda Red homologous recombination (61). The resulting strain was used as a donor for phage P1 transduction (62) to move the Spec^R^ marker and the linked *ara* operon into BW25113 to create the reference strain. The plasmid integration strain (Int2) was constructed by P1 transduction of the single-copy integration of the plasmid in the chromosome of strain B251 into BW25113. The *gfp* deletion strain (Del6) was made by transforming a plasmid from an evolved strain isolated from population 6 at day 20 into BW25113. Strains carrying both the pQGS plasmid and a satellite plasmid (Sat2 and Sat6) were made by transforming the purified SPs from strains B253 and B653, respectively, into the ancestor BW25113 / pQGS association. When constructing the SP strains, all cells in the initial transformation reaction were cultured through two growth cycles in LB-Spec to enrich for cells that acquired the SP. Then, these populations were plated on LB-Spec agar and colonies were screened for the presence of the SP by PCR and Sanger sequencing to identify the clones used in the fitness assays.

For each competition experiment comparing two strains, six colonies of the strain to be tested and six of the reference strain were picked to initiate separate overnight test tube cultures in 3 ml of LB (for the BW25113 versus reference strain competition) or LB-Spec (for all other competitions). These cultures were diluted 2000-fold into 10 ml fresh medium in 50 ml Erlenmeyer flasks and incubated for 24 h. These pre-conditioned cultures of each strain of interest were mixed with the reference strain in triplicate at volume ratios ranging from 1:1 to 28:1 depending on relative fitness of the two strains, while always maintaining an overall culture dilution of 2000-fold, into 10 ml of fresh medium to begin the competition. The ratio of each strain before and after 24 h of growth was determined by plating dilutions on tetrazolium arabinose (TA) agar plates. Relative fitness values were calculated for each of the total of 18 replicate competition assays for a pair of strains as previously described (63).

### Bee gut symbiont experiments

Plasmid pQGS was conjugated into bee gut symbiont *S. alvi* strain wkB2 as described previously (58). For the *in vitro* evolution experiment, five populations were started from single colonies that were inoculated into 6 ml of Insectagro DS2 medium (Corning) amended with 30 μg/ml Spec (DS2-Spec). *S. alvi* was cultivated in 6 ml of medium in culture tubes under 5% CO2 at 35°C without shaking. At 4-day intervals, 6 μl of each culture was transferred into 6 ml of fresh media and checked for the presence of SPs by PCR and Sanger sequencing.

For the *in vivo* evolution experiment, microbiota-free bees were obtained as described by Zheng *et al.* (44) with the following modifications. Late-stage pupae were removed manually from brood frames and placed in sterile plastic bins. The pupae emerged in an incubator at 35°C and humidity of 75%. Newly emerged bees were kept in cup cages with sterilized sucrose syrup (0.5 M) and bee bread. For inoculating these bees, *S. alvi* wkB2 cells containing the pQGS plasmid cultured on DS2-Spec agar were first suspended in 1×PBS to an OD600 of 1.0. Then, batches of 20-25 bees were placed into a 50-ml conical tube, and 50 μl sucrose syrup was added. The tube was rotated gently to coat the syrup on the surface of bees. Finally, 50 μl of the *S. alvi* wkB2/pQGS suspension (~5×10^7^cells) was added to the tube, and it was rotated again. *S. alvi* from the body surface colonizes the guts of these bees as a result of auto- and allogrooming.

Inoculated bees were reared in cup cages. Five of these enclosures were set up in each of two different experiments with more than 12 bees in each cage. The inoculated bees were fed sucrose syrup with spectinomycin (60 μg/ml) throughout the experiment. Bee guts were dissected 24 h after the inoculation, and wkB2/pQGS colonization was examined by fluorescence microscopy. To characterize SP evolution, one bee gut from each cup was dissected every day until 4 days after inoculation. Colonization levels were determined at day 3 and day 4 as colony-forming units (CFUs) present in dissected guts, as described by Kwong et al. (64). Bee guts were homogenized in 728 μl of cetyltrimethylammonium bromide (CTAB) buffer with 20 μl Proteinase K solution (20 mg/ml). DNA was then extracted using a bead-beating method as previously described (65). SPs were detected by PCR and characterized by Sanger sequencing. Repeat analysis

We created a Python script to enumerate microhomologies (short, near-perfect repeats) in pQGS that could mediate deletions leading to satellite plasmid formation or accessory-gene loss. We ran this code with settings that identified repeats that were at least 7 bases long, included at most one inserted or deleted base (indel), had at most five total mismatches (including indels), and with at least 75% sequence identity. When multiple valid alignments overlapped, only one was counted, giving precedence to the longer one, and then to the one with fewer mismatches.

### Plasmid evolution simulations

We implemented multilevel stochastic simulations that replicate cells and plasmids using Monte Carlo methods in Python. We established parameters that fit the observed numbers and types of plasmids typical for each cell type to its fitness by first assuming a linear cost for expression of each protein from the plasmid (66, 67). With 18 total plasmids per cell, an additive fitness model with a cost of 3.06% per full-length plasmid and 0.50% per deletion plasmid in a cell matches the experimentally observed fitness values for cells containing solely ancestral or solely deletion plasmids. A model in which satellite plasmids have no fitness cost and the collection of plasmids in each cell is exactly doubled and randomly but evenly assorted to daughter cells leads to an equilibrium level of 5.2 ancestral plasmids and 12.8 satellite plasmids per cell, on average. This is close to the ratio of the two types observed for several evolved strains. Note that instead of the expected relative fitness value of 0.841 for the cost of those 5.2 full-length plasmids, loss of all ancestral plasmids from some offspring due to segregation causes this population of cells to realize a relative fitness of just 0.77, which also closely matches the experiments. Alternative plasmid replication and segregation models that introduced more complexity and randomness gave lower fitness values and/or prevalence of satellite plasmids in cells than we observed. For example, we tested allowing plasmids to replicate via their own stochastic growth process and unevenly segregating plasmids between daughter cells.

The stochastic simulations track a collection of cells of interest that may contain different numbers of each plasmid type as they compete against a background of cells with a fixed fitness value and the cell population is transferred via serial dilution. We used them in three ways: (1) To determine the equilibrium fitness value of each cell type and equilibrium number of satellite plasmids per cell, we tuned the fitness of the background population in the model until a population seeded with 10,000 initial cells of the type of interest were able to maintain a stable population size over 500 generations of growth with 2-fold serial dilutions. (2) We examined the phenotypic delay of different plasmid mutations by starting 200 simulated populations with 20,000 cells containing 1 copy of either a satellite plasmid or a deletion plasmid and 17 copies of the ancestral plasmid and propagating them for 50 generations with 1.072-fold dilutions in a background population of cells containing only the ancestor plasmid. The realized fitness of mutant subpopulation at a given timepoint was calculated from the number of doublings it achieved relative to the background population within this time period. (3) To measure the chance that a new cell with a satellite plasmid or deletion plasmid would successfully escape dilution and establish under the conditions of our evolution experiment, we started 1,000,000 simulations, each with one cell containing one mutant plasmid that arose at a random cell division within a daily growth cycle with a 2,000-fold transfer dilution. Then, we followed the cell number of this population competing against a background of cells with only the ancestor plasmid until cells with the mutant plasmid either went extinct or reached a population size that was 20-fold higher than the dilution factor at the end of a growth cycle. The chance that a new mutant will establish is number of events of the latter type divided by the total number of trials.

### Code Availability

Python and R scripts for the repeat analysis and population genetics simulations are freely available online (https://github.com/barricklab/satellite-plasmid).

## Acknowledgments

We thank Nancy Moran for the use of her laboratory’s facilities for honey bee experiments. Sean Leonard and other researchers in the Barrick and Moran labs provided helpful feedback on this project. The authors acknowledge the Texas Advanced Computing Center (TACC) for providing high performance computing resources.

## References

1. Gogarten JP, Townsend JP (2005) Horizontal gene transfer, genome innovation and evolution. Nat Rev Microbiol 3:679–687.

2. Norman A, Hansen LH, Sørensen SJ (2009) Conjugative plasmids: vessels of the communal gene pool. Philos Trans R Soc Lond B Biol Sci 364:2275–2289.

3. Thomas CM, Nielsen KM (2005) Mechanisms of, and barriers to, horizontal gene transfer between bacteria. Nat Rev Microbiol 3:711–721.

4. Bryan LE, Van Den Elzen HM, Tseng JT (1972) Transferable drug resistance in *Pseudomonas aeruginosa*. Antimicrob Agents Chemother 1:22–29.

5. Wolters B, Kyselková M, Krögerrecklenfort E, Kreuzig R, Smalla K (2014) Transferable antibiotic resistance plasmids from biogas plant digestates often belong to the IncP-1ε subgroup. Front Microbiol 5:765.

6. Andersson DI, Hughes D (2010) Antibiotic resistance and its cost: is it possible to reverse resistance? Nat Rev Microbiol 8:260–271.

7. San Millan A, Toll-Riera M, Qi Q, MacLean RC (2015) Interactions between horizontally acquired genes create a fitness cost in *Pseudomonas aeruginosa*. Nat Commun 6:6845.

8. Bottery MJ, Wood AJ, Brockhurst MA (2017) Adaptive modulation of antibiotic resistance through intragenomic coevolution. Nat Ecol Evol 1:1364–1369.

9. San Millan A, MacLean RC (2017) Fitness costs of plasmids: a limit to plasmid transmission. Microbiol Spectr 5:1–12.

10. Hall JPJ, Wood AJ, Harrison E, Brockhurst MA (2016) Source–sink plasmid transfer dynamics maintain gene mobility in soil bacterial communities. Proc Natl Acad Sci 113:8260–8265.

11. Hall JPJ, Brockhurst MA, Dytham C, Harrison E (2017) The evolution of plasmid stability: Are infectious transmission and compensatory evolution competing evolutionary trajectories? Plasmid 91:90–95.

12. Bouma JE, Lenski RE (1988) Evolution of a bacteria/plasmid association. Nature 335:351–352.

13. Harrison E, Guymer D, Spiers AJ, Paterson S, Brockhurst MA (2015) Parallel compensatory evolution stabilizes plasmids across the parasitism-mutualism continuum. Curr Biol 25:2034–2039.

14. Loftie-Eaton W et al. (2017) Compensatory mutations improve general permissiveness to antibiotic resistance plasmids. Nat Ecol Evol 1:1354–1363.

15. Modi RI, Wilke CM, Rosenzweig RF, Adams J (1991) Plasmid macro-evolution: selection of deletions during adaptation in a nutrient-limited environment. Genetica 84:195–202.

16. Dahlberg C, Chao L (2003) Amelioration of the cost of conjugative plasmid carriage in *Eschericha coli* K12. Genetics 165:1641–1649.

17. Millan AS et al. (2014) Positive selection and compensatory adaptation interact to stabilize non-transmissible plasmids. Nat Commun 5:5208.

18. Harrison E, Brockhurst MA (2012) Plasmid-mediated horizontal gene transfer is a coevolutionary process. Trends Microbiol 20:262–267.

19. Stalder T et al. (2017) Emerging patterns of plasmid-host coevolution that stabilize antibiotic resistance. Sci Rep 7:4853.

20. Oliva M et al. (2017) A novel group of IncQ1 plasmids conferring multidrug resistance. Plasmid 89:22–26.

21. Poirel L et al. (2010) A novel IncQ plasmid type harbouring a class 3 integron from *Escherichia coli*. J Antimicrob Chemother 65:1594–8.

22. Yau S, Liu X, Djordjevic SP, Hall RM (2010) RSF1010-like plasmids in Australian *Salmonella enterica* serovar *Typhimurium* and origin of their *sul2-strA-strB* antibiotic resistance gene cluster. Microb Drug Resist 16:249–52.

23. Wen Y, Pu X, Zheng W, Hu G (2016) High prevalence of plasmid-mediated quinolone resistance and IncQ plasmids carrying *qnrS2* gene in bacteria from rivers near hospitals and aquaculture in China. PLoS One 11:e0159418.

24. Loftie-Eaton W, Rawlings DE (2012) Diversity, biology and evolution of IncQ-family plasmids. Plasmid 67:15–34.

25. Guerry P, van Embden J, Falkow S (1974) Molecular nature of two nonconjugative plasmids carrying drug resistance genes. J Bacteriol 117:619–630.

26. Meyer R, Hinds M, Brasch M (1982) Properties of R1162, a broad-host-range, high-copy-number plasmid. J Bacteriol 150:552–562.

27. Meyer R (2009) Replication and conjugative mobilization of broad host-range IncQ plasmids. Plasmid 62:57–70.

28. Sakai H, Komano T (1996) DNA replication of IncQ broad-host-range plasmids in gram-negative bacteria. Biosci Biotechnol Biochem 60:377–82.

29. Kuzminov A (2001) Single-strand interruptions in replicating chromosomes cause double-strand breaks. Proc Natl Acad Sci 98:8241–8246.

30. Rawlings DE, Tietze E (2001) Comparative biology of IncQ and IncQ-like plasmids. Microbiol Mol Biol Rev 65:481–496.

31. Chen L, Chen Y, Wood DW, Nester EW (2002) A new type IV secretion system promotes conjugal transfer in *Agrobacterium tumefaciens*. J Bacteriol 184:4838–4845.

32. Smalla K et al. (2000) Exogenous isolation of antibiotic resistance plasmids from piggery manure slurries reveals a high prevalence and diversity of IncQ-like plasmids. Appl Environ Microbiol 66:4854–62.

33. Clennel A, Johnston B, Rawlings DE (1995) Structure and function of Tn*5467*, a Tn*21*-like transposon located on the *Thiobacillus ferrooxidans* broad-host-range plasmid pTF-FC2. Appl Environ Microbiol 61:4223–4229.

34. Vogwill T, MacLean RC (2015) The genetic basis of the fitness costs of antimicrobial resistance: a meta-analysis approach. Evol Appl 8:284–295.

35. Morales VM, Bäckman A, Bagdasarian M (1991) A series of wide-host-range low-copy-number vectors that allow direct screening for recombinants. Gene 97:39–47.

36. Baba T et al. (2006) Construction of *Escherichia coli* K-12 in-frame, single-gene knockout mutants: the Keio collection. Mol Syst Biol 2:2006.0008.

37. Taton A et al. (2014) Broad-host-range vector system for synthetic biology and biotechnology in cyanobacteria. Nucleic Acids Res 42:e136.

38. Lovett ST, Hurley RL, Sutera VA, Aubuchon RH, Lebedeva MA (2002) Crossing over between regions of limited homology in *Escherichia coli:* RecA-dependent and RecA-independent pathways. Genetics 160:851–859.

39. Lee H, Doak TG, Popodi E, Foster PL, Tang H (2016) Insertion sequence-caused large-scale rearrangements in the genome of *Escherichia coli*. Nucleic Acids Res 44:7109–7119.

40. Chong P, Hui I, Loo T, Gillam S (1985) Structural analysis of a new GC-specific insertion element IS186. FEBS Lett 192:47–52.

41. saiSree L, Reddy M, Gowrishankar J (2001) IS*186* insertion at a hot spot in the *lon* promoter as a basis for Lon protease deficiency of *Escherichia coli* B: identification of a consensus target sequence for IS*186* transposition. J Bacteriol 183:6943–6.

42. Kwong WK, Moran NA (2013) Cultivation and characterization of the gut symbionts of honey bees and bumble bees: description of *Snodgrassella alvi* gen. nov., sp. nov., a member of the family *Neisseriaceae* of the *Betaproteobacteria*, and *Gilliamella apicola*. Int J Syst Evol Microbiol 63:2008–2018.

43. Kwong WK, Moran NA (2016) Gut microbial communities of social bees. Nat Rev Microbiol 14:374–384.

44. Zheng H, Powell JE, Steele MI, Dietrich C, Moran NA (2017) Honeybee gut microbiota promotes host weight gain via bacterial metabolism and hormonal signaling. Proc Natl Acad Sci 114:4775–4780.

45. Martinson VG, Moy J, Moran NA (2012) Establishment of characteristic gut bacteria during development of the honeybee worker. Appl Environ Microbiol 78:2830–2840.

46. Powell JE, Martinson VG, Urban-Mead K, Moran NA (2014) Routes of acquisition of the gut microbiota of the honey bee *Apis mellifera*. Appl Environ Microbiol 80:7378–7387.

47. Ilhan J et al. (2019) Segregational drift and the interplay between plasmid copy number and evolvability. Mol Biol Evol 36:472–486.

48. Rodriguez-Beltran J et al. (2018) Multicopy plasmids allow bacteria to escape from fitness trade-offs during evolutionary innovation. Nat Ecol Evol 2:873–881.

49. Haring V et al. (1985) Protein RepC is involved in copy number control of the broad host range plasmid RSF1010. Proc Natl Acad Sci U S A 82:6090–6094.

50. Frey J, Bagdasarian MM, Bagdasarian M (1992) Replication and copy number control of the broad-host-range plasmid RSF1010. Gene 113:101–106.

51. Tanaka K et al. (1994) Functional difference between the two oppositely oriented priming signals essential for the initiation of the broad host-range plasmid RSF1010 DNA replication. Nucleic Acids Res 22:767–772.

52. Becker EC, Meyer RJ (2003) Relaxed specificity of the R1162 nickase: A model for evolution of a system for conjugative mobilization of plasmids. J Bacteriol 185:3538–3546.

53. Kok M, Arnberg AC, Witholt B (1989) Single-stranded circular DNA generated from broad host range plasmid R1162 and its derivatives. Plasmid 21:238–241.

54. Swingle B et al. (2010) Oligonucleotide recombination in Gram-negative bacteria. Mol Microbiol 75:138–148.

55. Ithurbide S et al. (2015) Single strand annealing plays a major role in RecA-independent recombination between repeated sequences in the radioresistant *Deinococcus radiodurans* bacterium. PLoS Genet 11:e1005636.

56. Dutra BE, Sutera VA, Lovett ST (2007) RecA-independent recombination is efficient but limited by exonucleases. Proc Natl Acad Sci 104:216–221.

57. Pilla G, McVicker G, Tang CM (2017) Genetic plasticity of the *Shigella* virulence plasmid is mediated by intra- and inter-molecular events between insertion sequences. PLoS Genet 13:e1007014.

58. Leonard SP et al. (2018) Genetic engineering of bee gut microbiome bacteria with a toolkit for modular assembly of broad-host-range plasmids. ACS Synth Biol 7:1279–1290.

59. Deatherage DE, Barrick JE (2014) Identification of mutations in laboratory-evolved microbes from next-generation sequencing data using breseq. Methods Mol Biol 1151:165–188.

60. Deatherage DE, Traverse CC, Wolf LN, Barrick JE (2014) Detecting rare structural variation in evolving microbial populations from new sequence junctions using breseq. Front Genet 5:468.

61. Serra-Moreno R, Acosta S, Hernalsteens J, Jofre J, Muniesa M (2006) Use of the lambda Red recombinase system to produce recombinant prophages carrying antibiotic resistance genes. BMC Mol Biol 7:31.

62. Thomason LC, Costantino N, Court DL (2007) *E. coli* genome manipulation by P1 transduction. Curr Protoc Mol Biol Chapter 1:Unit 1.17.

63. Wiser MJ, Lenski RE (2015) A comparison of methods to measure fitness in *Escherichia coli*. PLoS One 10:e0126210.

64. Kwong WK, Engel P, Koch H, Moran NA (2014) Genomics and host specialization of honey bee and bumble bee gut symbionts. Proc Natl Acad Sci U S A 111:11509–11514.

65. Powell JE, Leonard SP, Kwong WK, Engel P, Moran NA (2016) Genome-wide screen identifies host colonization determinants in a bacterial gut symbiont. Proc Natl Acad Sci U S A 113:13887–13892.

66. Scott M, Gunderson CW, Mateescu EM, Zhang Z, Hwa T (2010) Interdependence of cell growth and gene expression: origins and consequences. Science 330:1099–1102.

67. Ceroni F, Algar R, Stan G-B, Ellis T (2015) Quantifying cellular capacity identifies gene expression designs with reduced burden. Nat Methods 12:415–418.

