## Supplemental Figures and Tables for "Evolution of satellite plasmids can stabilize the maintenance of newly acquired accessory genes in bacteria"

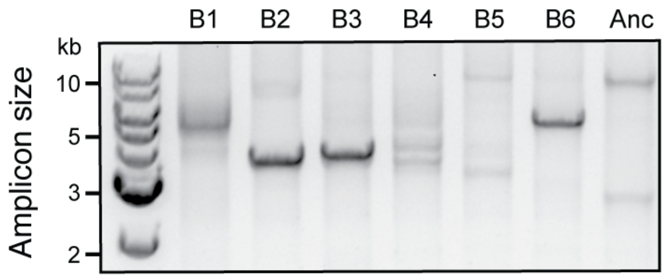


**Fig. S1. Satellite plasmids evolved in all *E. coli* populations.** Plasmid DNA was isolated from all six populations (B1–B6) of the *E. coli* evolution experiment on day 5. Then the PCR assay described in **Figure 1** was used to generate linearized amplicons to detect fragments with reduced sizes indicating that satellite plasmids were present in these genetically heterogenous populations.

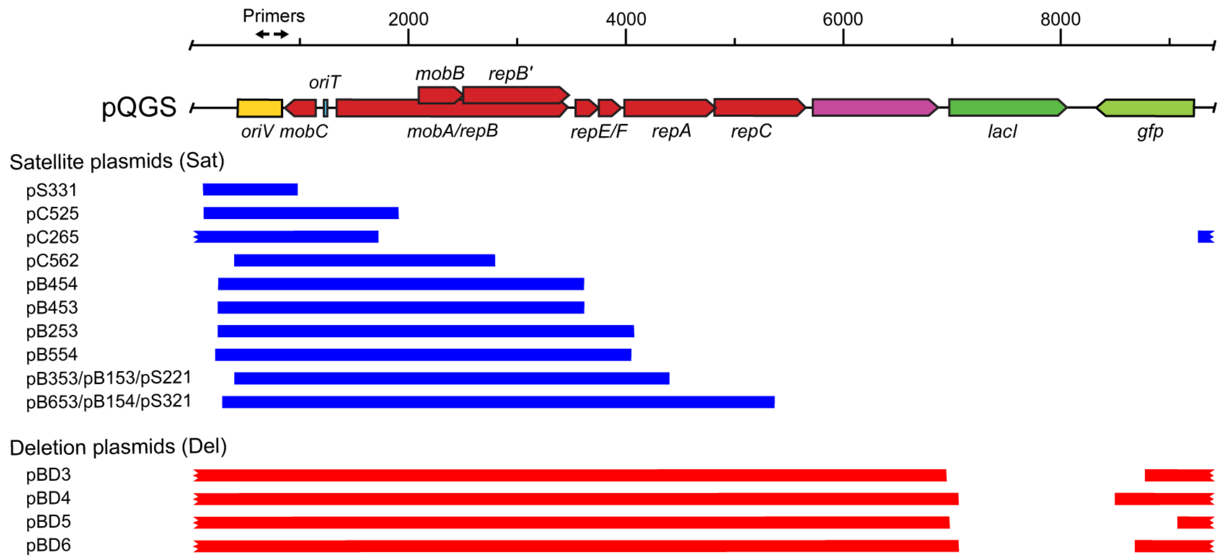


**Fig. S2. Maps of evolved plasmids found in all experiments.** Bars show regions preserved in plasmids that evolved in different *E. coli* (pB) and *S. alvi* (pS) populations cultured in the laboratory or in *S. alvi* populations that colonized the guts of honey bees reared in different laboratory enclosures (pC). The plasmid naming scheme is described in the legend to **Table S1**.

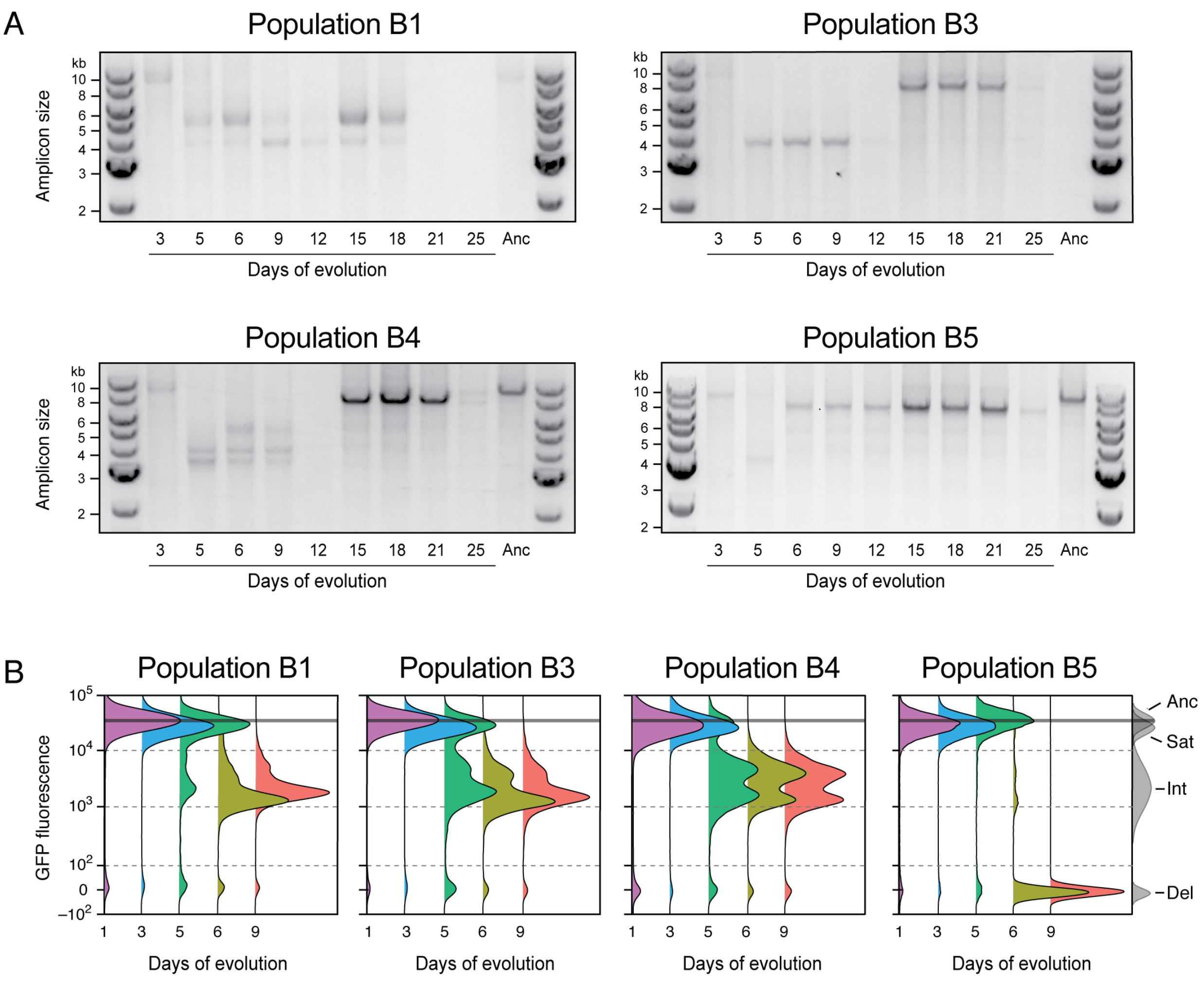


**Fig. S3. Time courses of plasmid evolution in the remaining *E. coli* populations.** (A) Appearance and persistence in populations B1, B3, B4, and B5 of satellite plasmids and plasmids with accessory-gene deletions that also reduce the size of the linearized plasmid PCR amplicon. (B) Distributions of GFP expression in cells in each population determined using flow cytometry.

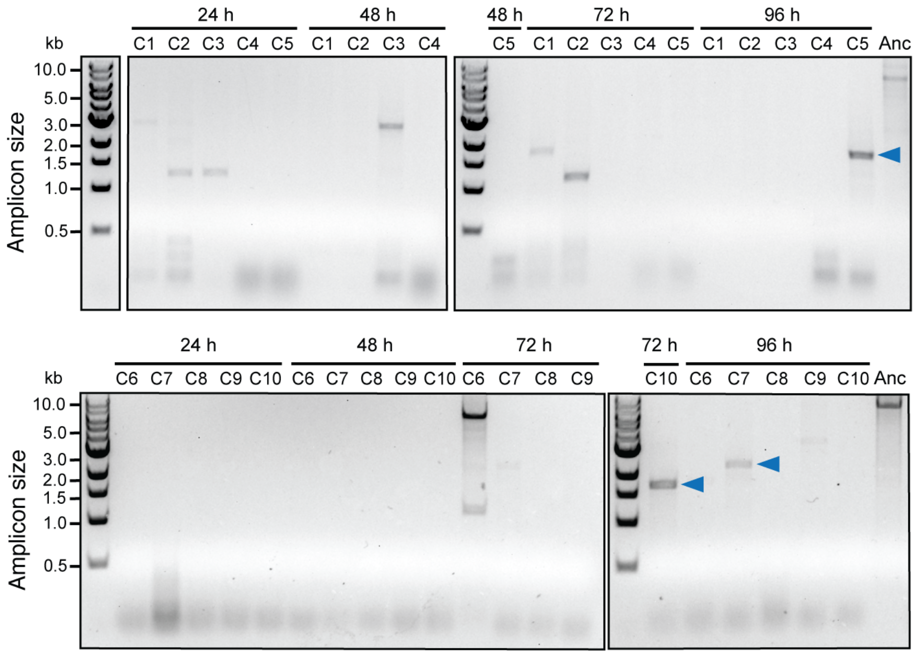


**Fig. S4. Evolution of satellite plasmids in *S. alvi* populations colonizing honey bee guts.** Multiple bees were reared in a total of ten separate enclosures in two sets of experiments conducted at different times. One bee from each enclosure (C1–C5 or C6–C10) was sacrificed at each of the indicated time points. DNA was isolated from each bee gut and subjected to the PCR assay to detect linearized amplicons from plasmids with deletions. Blue arrows in all panels are bands that were Sanger sequenced to validate that they represent satellite plasmids (**Fig. S2** and **Table S1**).

**Table S1. Sequence context of pQGS deletions forming satellite or deletion plasmids.**

| Evolved plasmid | Plasmid size (bp) | Endpoint coordinates | Flanking sequences (microhomology) |
|---|---|---|---|
| **Satellite plasmids (Sat)** | | | |
| pS331 | 854 | 964 | GATCCTCCgGCCaCt \| CGCTGCTGTTCACCT |
| | | 111 | CATGCTCCaGCCgCc \| CGCATTGGAGAAATT |
| pC1025 | 1793 | 1900 | GCGGGCTGGCCACGA \| CGCCCGCATTGACCA |
| | | 108 | AAGCCATGCTCCAGC \| CGCCCGCATTGGAGA |
| pC765 | 1863 | 1708 | CCCGCACTGCCACcT \| GATGATCTCCGAGCG |
| | | 9243 | GCGACGAATTGCCAC \| gTTGTCGCAGTGTCT |
| pC562 | 2396 | 2783 | GGCCGGGAGCCTGCC \| CTGGTAGTGGAAACC |
| | | 388 | GCCGAAATGCCTGCC \| GTTGCTAGACATTGC |
| pB454 | 3359 | 3606 | GGTTGCCCGGTGGCT \| tTGGTTATACGTCAA |
| | | 248 | CTCGGGTCGGTGGCT \| cTGGTAACGACCAGT |
| pB453 | 3360 | 3606 | GGTTGCCCGGTGGCT \| tTGGTTATACGTCAA |
| | | 247 | GCTCGGGTCGGTGGC \| TcTGGTAACGACCAG |
| pB253 | 3835 | 4081 | CGGTACGGTCGGGGC \| gCTGGTGTCGCCCGG |
| | | 247 | GCTCGGGTCGGtGGC \| tCTGGTAACGACCAG |
| pB554 | 3836 | 4072 | CATGGTGGCCGGTAC \| GGTCGGGGCgCTGGT |
| | | 237 | AGAAGGGTTTGCTCG \| GGTCGGtGGCtCTGG |
| pB353 / pB153 / pS221 | 4003 | 4401 | TGATGGTGCTgGACA \| CGCTGCGCCGGTTCC |
| | | 399 | TGCCGTTGCTaGACA \| TTGCCAGCCAGTGCC |
| pB653 / pB154 / pS321 | 5077 | 5361 | ATCGCGCAGGCCGTC \| aTGGGTGGCGGCCAG |
| | | 285 | TCCCGGCTGGCCGTC \| cTGGCCGCCACATGA |
| **Deletion plasmids (Del)** | | | |
| pBD3 | 7534 | 6912 | TAGGATACAGAAACA \| GAGGAGATATTACGG |
| | | 8776 | TTAACAAGGGTATCA \| CCCTCGAACTTCACT |
| pBD4 | 7938 | 7035 | TCCCGCGTGGTGAAC \| CAGGCCAGCCACGTT |
| | | 8495 | GCTTTTCGTTGGGAT \| CTTTCGAAAGGGCAG |
| pBD5 | 7267 | 6942 | TACGCAAGTACACAA \| GATACAGGAGAGGTA |
| | | 9073 | ACATCACCGTCTAAT \| TCCACGAGGATTGGG |
| pBD6 | 7744 | 7024 | ATCAGACCGTTTCCC \| GCGTGGTGAACCAGG |
| | | 8678 | CCATAATGTACACAT \| TATGGGAGTTATAGT |

Evolved satellite plasmids are named as follows: with a letter for the experiment in which they were observed (pB, *in vitro E. coli*; pS, *in vitro S. alvi*; pC, *in vivo S. alvi*); then the index of the experimental population in which they were observed (1–10), then a digit for the transfer or time point at which they were isolated, and with a final digit representing a distinct clonal isolate from that population or sequenced PCR amplicon from that sample. Evolved deletion plasmids were all from the *E. coli* experiment and are labeled pBD with single unique index. Endpoint coordinates are for the terminal bases remaining in the pQGS sequence (Genbank:MH423581) that flank the deleted portion of each plasmid. The flanking sequences have a vertical bar showing the location where the deletion begins in the upper row and ends in the lower row. The portions of these flanking sequences that are shaded grey are deleted in evolved plasmids. Therefore, the sequence of the new plasmid maintains the bases on the left side of the upper row and then continues into the bases on the right side of the lower row. Microhomologies that appear to mediate the formation of satellite plasmids are highlighted in red with lowercase letters for mismatched bases.
